# Gambling on an empty stomach: Hunger modulates preferences for learned but not described risks

**DOI:** 10.1101/2021.03.19.435837

**Authors:** Maaike M.H. van Swieten, Rafal Bogacz, Sanjay G. Manohar

## Abstract

We assess risks differently when they are explicitly described, compared to when we learn directly from experience, suggesting dissociable decision-making systems. Our needs, such as hunger, could globally affect our risk preferences, but do they affect described and learned risks equally? On one hand, explicit decision-making is often considered flexible and context-sensitive, and might therefore be modulated by metabolic needs. On the other hand, implicit preferences learned through reinforcement might be more strongly coupled to biological drives. To answer this, we asked participants to choose between two options with different risks, where the probabilities of monetary outcomes were either described or learned. In agreement with previous studies, rewarding contexts induced risk-aversion when risks were explicitly described, but risk-seeking when they were learned through experience. Crucially, hunger attenuated these contextual biases, but only for learned risks. The results suggest that our metabolic state determines risk-taking biases when we lack explicit descriptions.

## Introduction

When we decide between options with uncertain outcomes, we factor risk into the decision. This is most commonly evaluated by asking people to decide between explicitly described, hypothetical choice scenarios (Allais, 1953; Arrow, 1951; Ellsberg, 1961; Kahneman and Tversky, 1979; Weber et al., 2004). In these experiments, risk-taking is typically modulated by the magnitude and probability of outcomes, or by framing choices in a positive or negative context using words or diagrams. This contrasts with real life scenarios, in which humans usually make repeated choices, and learn about uncertain outcomes from experience. Several studies have reported that experienced-based choices differ from choices based on verbal or graphical descriptions (Hertwig et al., 2004; Hertwig and Erev, 2009; Niv et al., 2012). This observation is better known as the experience-description gap. In particular, empirical studies have also shown that people are typically risk-seeking for negatively framed choices, but risk-averse for positively framed choices when outcomes are explicitly described (Kahneman and Tversky, 1979; Tversky and Kahneman, 1981). However, when experiential choices are framed in a positive or negative context, risk attitudes are reversed compared to description-based decisions (Hertwig et al., 2004; Ludvig et al., 2014; Ludvig and Spetch, 2011).

The effect of decision context is thought to be driven by anticipatory emotions (De Martino et al., 2006) as well as biological needs (Stephens, 1981). Nevertheless, only a handful of studies has investigated the effect of physiological factors, such as hunger, on explicit risk-taking behaviour in humans, and suggest that hunger increases risk-seeking (Levy et al., 2013; Shabat-Simon et al., 2018; Symmonds et al., 2010), but the effect of hunger on experiential risk-taking has not yet been tested in humans. Biological need, which is described as the disparity between the current state and the goal state, has been shown to motivate decision-making in animals that make experiential choices (Aw et al., 2011; Papageorgiou et al., 2016; Pompilio et al., 2006) and has been captured by computational models (van Swieten and Bogacz, 2020). The concept of making decisions to reduce this disparity also underlies the risk-sensitive foraging theory (Stephens, 1981). This theory describes that if the goal cannot be reached with a safe, low-risk option, then an individual should choose a high-risk option because it offers a chance of meeting the need and increases the chance of survival.

For described risks, the contextual modulation of risk-taking can be captured by a utility function, such as proposed by prospect theory (Kahneman and Tversky, 1979). For experienced risks, in contrast, contextual modulation can be accounted for by a recently described model that captures the roles of dopamine in learning and choice (Möller et al., 2021). This model aligns with evidence that dopamine enhancement promotes risk-seeking behaviour (Gallagher et al., 2007; Rigoli et al., 2016; St Onge and Floresco, 2009). The Prediction Error Induced Risk-Seeking (PEIRS) model proposes that a positive decision context elicits a positive context prediction error that enhances dopamine release, while a negative decision contexts evokes a negative context prediction error that reduces dopamine release (Möller et al., 2021). Similar to the utility function in prospect theory, PEIRS includes a risk sensitivity parameter that determines the impact of context on risk-taking. Crucially, if hunger alters the extent to which context modulates risk-taking, this could be captured by changes in this parameter.

Given that experiential and description-based risk-taking are thought to involve different neural systems (Fitzgerald et al., 2010), we tested two alternative hypotheses about the effects of hunger on explicitly described versus experientially learned risky choice. On one hand, we might expect the description-based decision-making to be modulated by hunger, because risk is tracked and represented in a range of cortical areas that are informed by high-level cognitive representations (Clark et al., 2008; Elliott et al., 1999; Hsu et al., 2005; Huettel et al., 2005, 2006; Knutson and Bossaerts, 2007; Kuhnen and Knutson, 2005; McCoy and Platt, 2005; O’Neill and Schultz, 2010; Platt and Huettel, 2008; Preuschoff et al., 2008; St. Onge et al., 2011; Tobler et al., 2007). It is susceptible to framing effects, whereby the cognitive, numerical and linguistic context of options influences choice (Allais, 1953; Arrow, 1951; Kahneman and Tversky, 1979) and might therefore be more flexible than the experienced-based system. Hunger may modulate high-level decision-making systems, with the appetite-stimulating hormone ghrelin activating receptors distributed widely in the cerebral cortex including hippocampus (Zigman et al., 2006) and can enhance memory and performance (Diano et al., 2006). Accordingly, hunger may increase risk-seeking for explicitly described food but also monetary reward (Levy et al., 2013; Shabat-Simon et al., 2018; Symmonds et al., 2010), suggesting that metabolic signals do impact cognitive decisions.

On the other hand, we might expect experiential decision-making to be biased by the organism’s needs, because it may rely more on primitive neural systems. The modulation of risk preferences according to energy reserves may be crucial for the adaptation to changes in the environment, in particular when resources are scarce (Houston, 1991; Kacelnik and Bateson, 1997; Stephens, 1981). Experiential decision-making relies on subcortical brain areas such as the striatum and the dopaminergic midbrain (Abler et al., 2006; Knutson et al., 2001; Niv et al., 2012; Tobler et al., 2007) that are targeted by circulating hormones that signal current energy reserves (Elmquist et al., 1998; Zigman et al., 2006). In particular, leptin inhibits and ghrelin activates dopaminergic neurons in the ventral tegmental area, and could therefore modulate learning and decision-making via the mesolimbic pathway (Abizaid et al., 2006; Figlewicz et al., 2007; Hommel et al., 2006). In line with this, in animal studies, food deprivation increases risk-seeking in experience-based tasks (Kacelnik and Bateson, 1997). Perhaps surprisingly, the effects of hunger on experiential and explicit risk-taking have never been directly compared.

We employed two complementary risk-taking tasks in a within-subject design. One task involved decisions between two options whose probability of winning and losing, and the magnitude of rewards, were explicitly described. The other task involved decisions between options whose average reward and uncertainty had to be learned through sampling.

We included three decision contexts to verify whether choices were driven by the expected value or by the risk of options. The options presented in a *mixed* context differed in their expected value, which typically drive risk-neutral behaviour. The pair of options in a *negative* context differed in risk, and were matched in expected value, but both yielded less than the average reward in the task. Options in the *positive* decision context were analogous to the negative context, but the expected values were both higher than average. These three decision contexts allowed us to examine the effect of both hunger and decision context on experiential and explicit risk-taking.

In agreement with previous studies, we showed that risk attitudes for described risks were opposite to those for learned risks. Hunger only modulated risk preferences for learned risks in a context-specific manner, showing that the experience-based system, but not the cognitive system, is sensitive to the motivational drive of an organism.

## Methods

### Participants

Thirty-two healthy volunteers (females: 20, mean age: 25.6 ±6.5) were recruited for this study. All participants were healthy, had no history of psychiatric diagnoses, neurological or metabolic illnesses, and had not used recreational drugs in the past 3 months. All participants had a normal weight (Body Mass Index: 22.9 ±3.2 kg/m^2^), regular eating patterns and no history of eating disorders. Each participant gave written informed consent and the study was conducted in accordance with the guidelines of the University of Oxford ethics committee. We estimated the effect size from previous papers as 0.25 (Shabat-Simon et al., 2018; Symmonds et al., 2010). We then used G*Power (3.1.9.7) and estimated that we need 30 participants to obtain a power of 0.85. *Post-hoc* power calculations confirmed that the observed power in our study was 0.8. All data and code is openly available at http://...

### Manipulation of metabolic state

Participants were tested in a within-subjects counterbalanced, randomised crossover design for the effects of food deprivation on risk-taking tasks (Fig. 1A). Sessions were approximately 1 week apart (at least 4 days, but no more than 14 days). All sessions took place at the same time of day between 10 am and 1 pm, to minimise time-of-day effects. For one session, participants were asked to refrain from eating and drinking caloric drinks from 8 pm the night prior to testing. For the other session, participants were asked to eat normally the day before and consume a full breakfast within 1 hour of arriving at the lab for testing. We assessed the effect of food deprivation on self-reported feelings of hunger and mood using a computerised Visual Analogue Scale of each session (Bond and Lader, 1974; Flint et al., 2000). Participants were asked to place a cursor on a 100 mm scale with positive or negative text ratings anchored at either end. This assessment provided a subjective measure of whether the manipulation worked. Participants performed the decision-by-description task first, then a learning, attention and planning not described in this paper, and finished with the decision-by-experience task. This order was fixed to control for fasting time. Finally, the session order did not affect performance.

**Figure 1:**
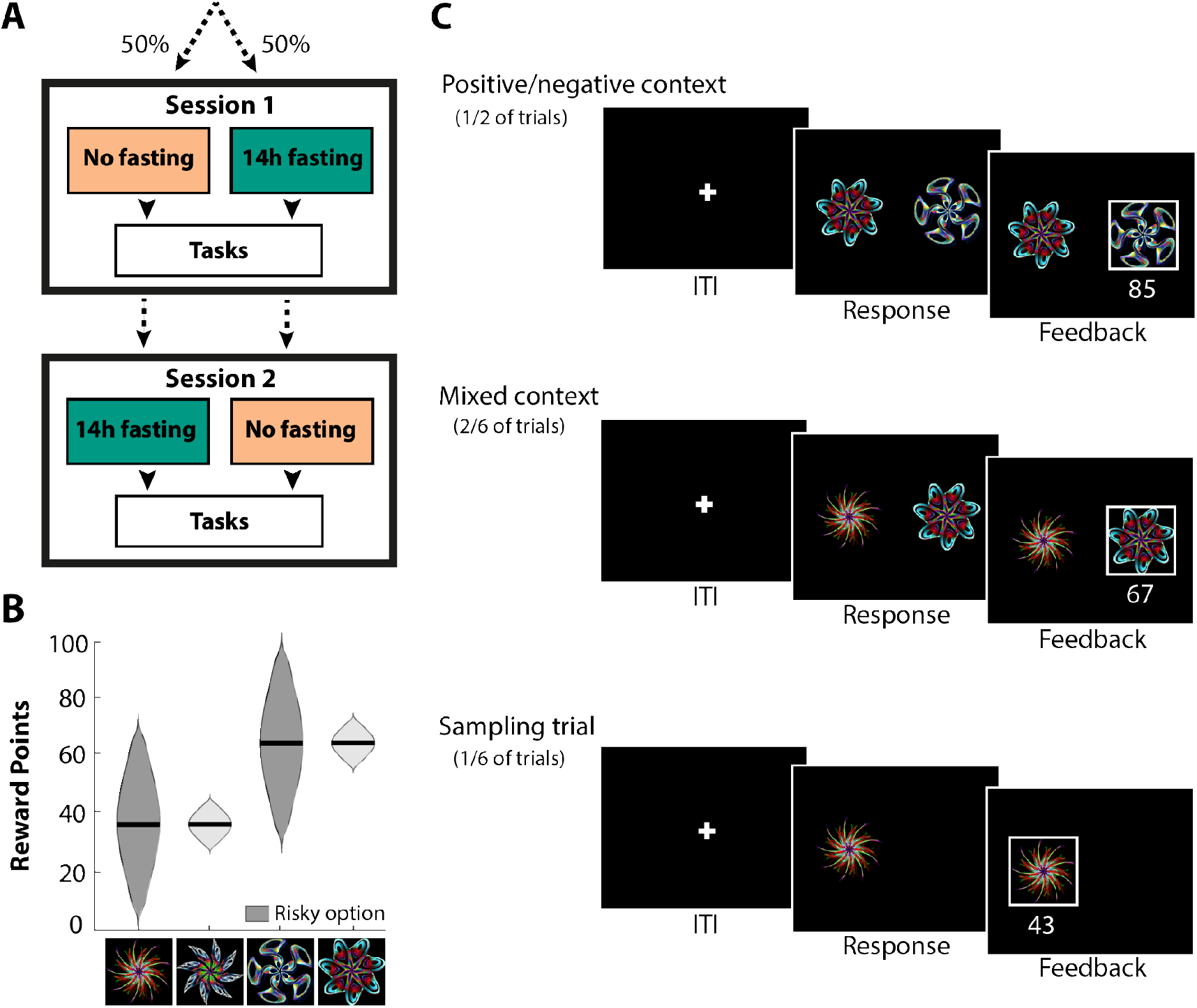
Decisions by experience. **A)** Participants were tested in a counterbalanced randomised crossover design. Participants were tested on two separate days approximately 1 week apart. One session took place after 14 hours fasting, the other session after consuming a full meal. **B)** Each reward distribution associated with a stimulus was approximately normal. The mean of the distribution was either 35 or 65 with a standard deviation of either 5 or 20. The dark grey distributions indicate the more risky option. **C)** Task structure of decisions made from experience. The task consisted of three different trial types: positive/negative context trials (1/2 of the trials), mixed context trials (2/3 of the trials) and sampling trials (1/6 of the trials). After a response, a reward sampled from the associated reward distribution was presented.

### Experimental design

#### Decisions by experience

We employed a modified version of a risk taking task developed by Möller et al. (2021). Participants learned the reward value of four stimuli through repeated sampling. Each stimulus was associated with a Gaussian reward distribution that followed a two-by-two design: high or low mean value (65 or 35 points) and high or low standard deviation (20 or 5) (Fig. 1B). When a stimulus was chosen, participants received a reward drawn from the corresponding distribution. The task included three trial types: positive/negative context trials (50%), mixed context trials (33%) and sampling trials (17%) (Fig. 1C) (Ludvig et al., 2014; Niv et al., 2012). Positive/negative context trials consisted of two options with equal mean, but different risks. Positive context trials have a mean above the average outcome in the task. In contrast, negative context trials have a mean below the average outcome in the task. Mixed context trials offered choices between options with unequal expected value, which were used to test whether participants paid attention to their choices and understood the difference between the stimuli. Sampling trials were forced choice trials in which only one stimulus was presented. These trials ensured that all options were sampled from and that participants occasionally experienced reward contingencies that they did not prefer (Ludvig et al., 2014; Niv et al., 2012).

Each trial had the same structure. After a short inter-trial-interval (ITI) of 500-700 milliseconds, the stimuli were presented on the screen. Responses were made by pressing on the left or right arrow key of the keyboard to choose the left or right option, respectively. Choices were immediately followed by feedback for 1.5 seconds, showing the number of points won (Fig. 1C). The total accumulated points was continuously displayed at the top of the screen. Participants were instructed to maximise their total number of points, which was converted into a monetary performance bonus at the end of the task. Each participants completed four blocks of 72 trials. All trial types were equally distributed over the blocks, but we ensured that a stimulus presented in a sampling trial did not precede a positive/negative context trial with the same stimulus to avoid priming of choices. Reward distributions were generated at the start of each block to ensure each block had the intended reward distribution and stimulus sets were reset after 2 blocks (or 144 trials). After each block, participants were asked to indicate the reward distribution of each stimulus by placing two cursors on a Visual Analogue Scale ranging from 0 to 100 points, one for the minimum and one for the maximum reward in the distribution. The rated spread was computed as the difference between the rated minimum and maximum of the reward distribution and the rated mean was taken as the average of the two values.

#### Decisions by description

Explicit risk-taking behaviour was probed using the probabilistic task described by Rogers and colleagues (Norbury et al., 2013; Rogers et al., 2003). On each trial, participants were required to choose between two simultaneously presented gambles (Fig. 2A). Each gamble was represented visually by a histogram of which the height indicated the relative probability of winning a given number of points. The magnitude of possible points to win was indicated in green above each histogram, with the magnitude of possible points to lose indicated below in red. We used 10 different types of choice (Fig. 2B), each specifying a decision between two options. Each of the two options was specified by a probability of winning, an amount that could be won, and an amount that could be lost. Eight of these 10 types of decision offered a choice between two options that differed in their objective expected values (mixed decision context). One option was a reference option, and always consisted of a 50:50 chance of winning or losing 10 points, giving an expected value of 0. The other option was a risky gamble, and varied either in the probability of winning (0.6 or 0.4), the magnitude of possible points to win (30 or 70 points), or the magnitude of possible points to lose (30 or 70 points). The remaining two decision types offered a choice between a certain win or loss and a 50:50 chance gamble with the *same expected value*. These gamble types were identical in terms of prospect, but differed in valence, allowing for the examination of positive and negative context effects on differences in risk attitudes. Visual feedback (win/lose) was given after each choice was made, and the revised running total points was presented before the next trial. Participants were instructed that each gamble should be considered independently of outcomes of previous gambles. Participants completed four blocks of 20 trials, and the order in which gambles were presented was kept constant for both conditions. The highest total score obtained in a block was converted into pence and paid at the end of the task as a performance bonus. Deliberation times were also recorded.

**Figure 2:**
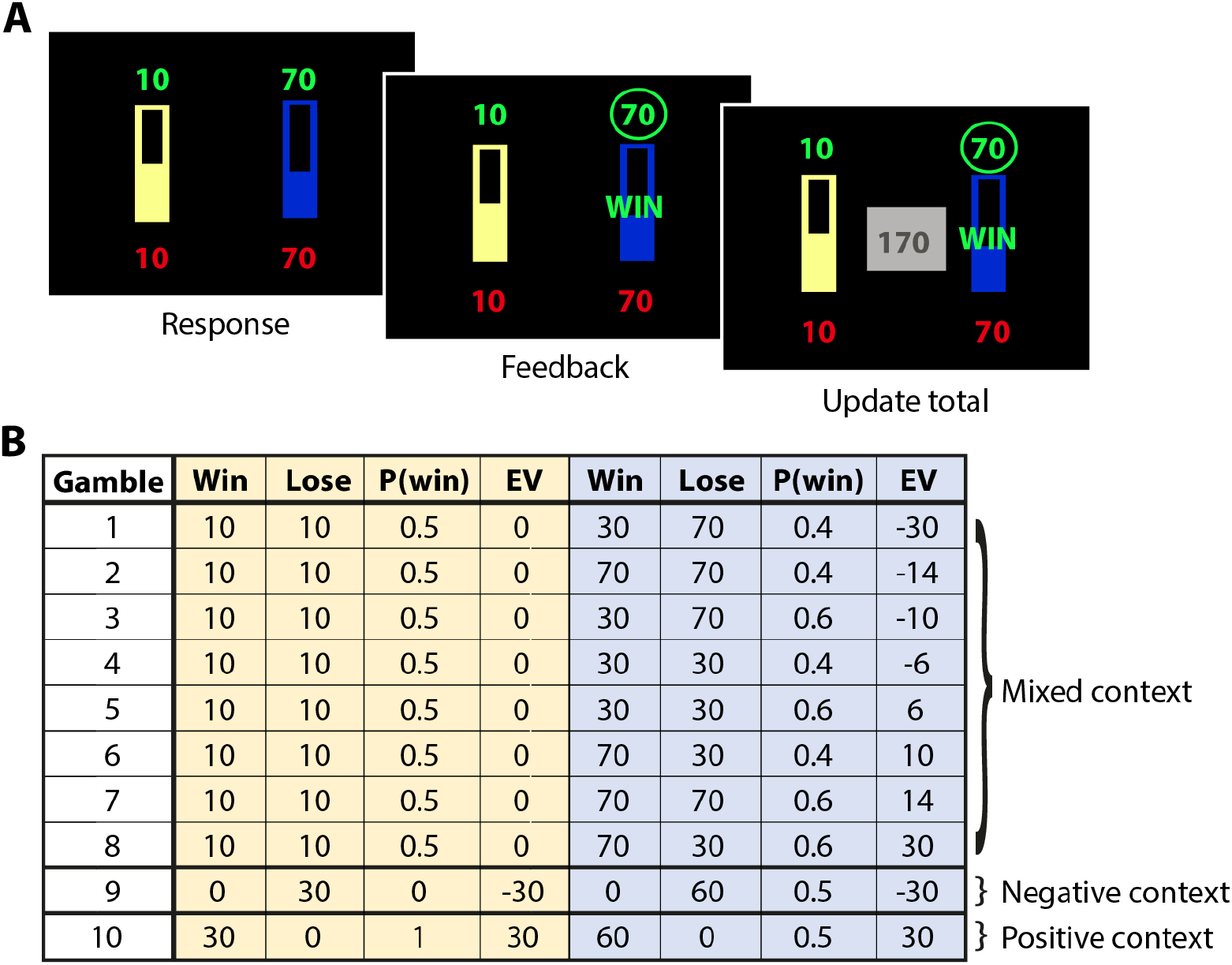
Decisions by description. **A)** Example trial of decisions from description. Each trial consisted of a choice between a reference gamble (yellow) and risky gamble (blue). Points to win and lose were presented in green and red, respectively. The probability of winning corresponded to the size of the filled bar. Feedback was given after a choice and the running total was updated. **B)** Gambles 1–8 show different combinations of points to win, points to lose and the probabilities of winning, with expected values ranging from −30 to 30. Gambles 9 and 10 have equal expected values, but different risks.

All computerised behavioural paradigms were implemented using Psychophysics Toolbox Version 3 on MATLAB (version 19b, MathWorks, Natick, MA).

### Behavioural analyses

Risk was defined as the uncertainty in possible outcomes of a decision, expressed as the variance of the associated reward distribution (Rothschild and Stiglitz, 1970). Risk attitudes were computed separately for positive and negative contexts.

For learned risks, the risk preference was averaged over the second half of the trials (72 trials) of each stimulus set (Fig. S1A). Using only the second half of the trials allowed participants sufficient opportunity to learn the outcomes associated with each option, while providing a long enough sample to get a reliable measure of their risk preference (Ludvig et al., 2014; Niv et al., 2012).

For described risks, the risk preference was assessed as the proportion of risky gambles chosen in the negative (decision type 9) or positive (decision type 10) decision context. All trials were included, because no learning occurred and each gamble was considered independently (Fig. S1B).

We used the performance on mixed context trials as a control measure to verify if people maximised their outcome. The proportion of options with the highest expected value was calculated based on the performance on mixed context trials in experiential risk-taking task and the gambles 1–8 in the explicit risk-taking task.

Statistical significance was tested using paired t-tests or repeated measures analysis of variance (ANOVA) as appropriate in MATLAB and SPSS (IBM Corp. Released 2019. IBM SPSS statistics for Windows, Version 26.0. Armonk, NY: IBM Corp.).

### Computational model fitting

We used two reinforcement learning models to further assess the effects of hunger on experience-based risk-taking. The models themselves are described in the results section. We used a hierarchical model-fitting strategy that takes into account the likelihood of individual participant choices given the individual participant parameters and also the likelihood of the individual participant parameters given the parameter distribution in the overall population across conditions. This two-stage hierarchical procedure is a estimation strategy of the iterative expectation-maximization algorithm (EM) (Guitart-Masip et al., 2011; Huys et al., 2012; MacKay, 2003). This regularises individual participants’ parameter fits, rendering them more robust toward over-fitting. To infer the maximum-a-posteriori estimate of each parameter for each participant, we set the prior distribution to the maximum-likelihood given the data of all participants and then use EM for the two conditions separately to obtain parameter estimates for each condition. The statistical significance was tested using paired t-tests with respect to the Gaussian scaled model parameters (see supplemental material for the transformation of parameters). Reported p-values were corrected for multiple comparisons using the Bonferonni method.

In the fitting procedure, all context trials were used to estimate all parameters. Sampling trials were only included for the estimation of learning rates for the mean and variance of a stimulus using Eq. (1) and Eq. (3), respectively. Due to the absence of a choice, sampling trials were excluded from the estimation of the softmax choice parameter and the risk parameters. The presence of only one stimulus makes the probability of choosing this stimulus one, and this would interfere with the parameter estimation. Initial values for *Q* and *S* were set to 50 and 5, respectively. The model comparison and parameter recovery method can be found in the supplemental material.

## Results

As expected, participants rated their subjective feelings of hunger significantly higher after 14 hours of fasting than after eating a full meal (Wilcoxon signed rank test: [*Z* = −4.84, *p* < 0.0001, *d* = 0.86]), indicating that the manipulation was successful.

### Hunger altered experiential risk-taking in a context-specific manner

We first analysed choice behaviour in the positive and negative context to evaluate experiential risk-taking in a context-specific manner (Fig. 3A). Participants were significantly more likely to choose the risky option in a positive decision context, but not a negative context (main effect of context [*F*_1,31_= 10.28, *p* < 0.003, 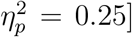]). Such risk-seeking for positive and risk avoidance for negative decision contexts is consistent with previously reported risk attitudes for learned risks (Ludvig et al., 2014; Madan et al., 2015). Crucially, food deprivation modulated risk-attitudes for positive and negative contexts in opposite manner (interaction effect of food deprivation and context [*F*_1,31_= 8.38, *p* < 0.007, 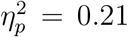]), such that hunger neutralised the risk preferences in both contexts. A *post hoc* paired t-test revealed that this interaction effect was mainly driven by hunger decreasing risk-taking behaviour in the positive context [*t*_31_ = 2.73, *p* = 0.010, *d* = 0.49], and not by an increase in risk-seeking in the negative context [*t*_31_ = 1.01, *p* = 0.319, *d* = 0.18].

**Figure 3:**
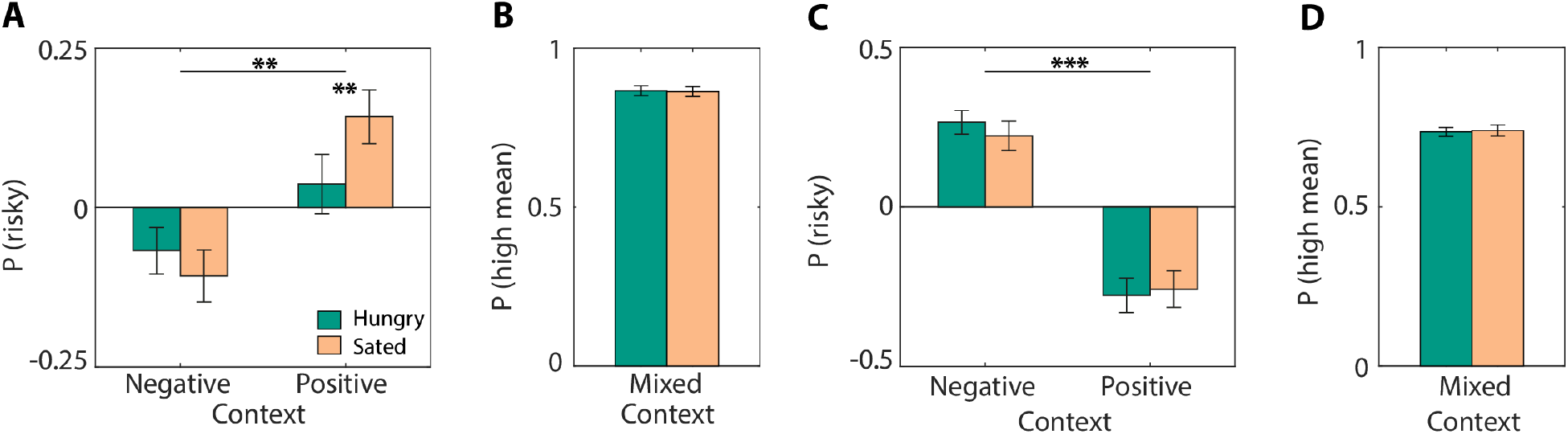
Risk attitudes for learned and described risks. **A)** For learned risks, participants were risk-averse for negative decision contexts and risk-seeking positive decision contexts. Food deprivation attenuated risk attitudes for decision contexts in opposite direction. **B)** Proportion of high mean options chosen for mixed context trials in decisions from experience. **C)** For described risks, participants were risk-seeking for negative contexts and risk-averse for positive contexts (gambles 9 and 10; Fig. 2B). Food deprivation did not affect these risk preferences. **D)** Proportion of high mean options chosen for mixed context trials (gambles 1–8; Fig. 2B) in explicit risk task. Error bars represent SEM. ** *p* < 0.01, *** *p* < 0.001.

Although the interaction is significant at a group level, we further asked whether the effect is strong enough to be seen within individuals. For each participant, we ran a *post hoc* context × hunger logistic regression (Fig. S2). 10 out of 32 people had effects that reached significance in the expected direction even within single participants. Only 1 person had a significant effect in the opposite direction. Finally, food deprivation did not alter overall risk-taking behaviour (main effect of food deprivation [*F*_1,31_ = 1.19, *p* = 0.283, 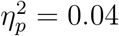]).

To verify that neutral risk preferences were not caused by an inability to differentiate stimuli, we used mixed context trials to examine whether participants understood the difference in mean and variance of reward distributions. All participants performed on average above 90% on mixed context trials, and no participant performed below 60%, indicating that they understood the distinction between high and low mean stimuli. The level of food deprivation did not affect the accuracy on mixed context trials [*t*_31_ = 0.62, *p* = 0.543; Fig. 3B].

Finally, the observed changes in risk preferences following food deprivation were not the result of changes in attention, as the overall reaction times were consistent across conditions (main effect of food deprivation [*F*_1,31_ < 1]; Fig. S3A).

### Hunger did not affect explicit risk-taking

To provide a comparable measure to the context effects in experience-based risk-taking, we also analysed the risk preference for matched mean gambles in positive and negative decision context in description-based choices (Fig. 3C). Participants were risk-seeking for negative decision contexts and risk-averse for positive decision contexts (main effect of context [*F*_1,31_ = 55.01, *p* < 0.0001, 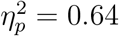]). This risk pattern has been previously described by prospect theory (Kahneman and Tversky, 1979), in which extreme positive outcomes are down-weighted. In contrast to learned risks, food deprivation did *not* alter context-specific risk attitudes for described risks (interaction effect of food deprivation and context [*F*_1,31_ = 1.53, *p* = 0.255, 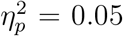]) or overall risk-taking (main effect of food deprivation [*F*_1,31_ < 1]; Fig. 3C). In line with previous reports, the risk attitudes for experiential and explicit risk-taking were opposite, and confirms the existence of the description-experience gap (Hertwig and Erev, 2009).

Participants chose the option with the highest expected value more often in mixed decision contexts, regardless of the level of risk (Fig. 3D), showing that the difference in expected value drove choice behaviour (Weber et al., 2004). In line with the performance on the experiential task, but inconsistent with previous findings (Levy et al., 2013; Symmonds et al., 2010), food deprivation did not attenuate this effect [*t*_31_ = −0.29, *p* = 0.776]. Despite the absence of a shift in risk preference, food deprivation increased reaction times for all gambles independently of the decision context (main effect of food deprivation [*F*_1,31_ = 37.42, *p* < 0.0001, 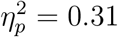]; Fig. S3B).

### Modelling of risk-sensitive choice behaviour

The previous analyses showed that food deprivation only altered decision-making when risks had to be learned. However, the behavioural analyses do not provide insight into what computational process was altered by food deprivation. Therefore, we employed a computational modelling strategy to account for the integration of a specific reward history triggered by sampling. This strategy allowed us to attribute the effects of food deprivation to a specific computational process. We relied on two reinforcement learning models to explain the behavioural data: a standard reinforcement learning (RW) model (Rescorla and Wagner, 1972) and an adapted version of a recently proposed reinforcement learning model that can account for contextual risk preferences (PEIRS) (Möller et al., 2021).

Standard reinforcement learning models provide trial-by-trial estimates of the expected mean value of each stimulus, without considering the variability in outcomes. They provide a good account of what a rational decision-maker would do based on the objective expected value of a reward. In this model, the expected mean value of the chosen stimulus *Q_c_* was updated using:

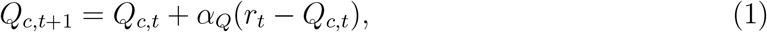

where *r_t_* is the reward obtained by choosing a stimulus on trial *t* and *α_Q_* is the learning rate for the mean reward. Decisions in this model were solely based on the expected mean value of the presented stimuli. The utility *U* of stimulus i on trial *t* was *U_i,t_* = *Q_i,t_*. The probability of choosing an option was computed using the softmax decision rule:

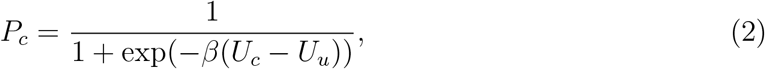

where *U* reflects the utility for the chosen *c* and unchosen *u* option. The parameter *β* determines the participant’s tendency to exploit (i.e. to choose the stimulus with the highest *U* value) or to explore (i.e. to randomly choose a stimulus).

In contrast, the PEIRS model accounts for both the average outcome and the variability, or spread (*S*), in outcomes of an action. It also captures innate risk propensities and assumes that positive and negative decision contexts influence how the spread in reward outcomes affects the subjective utility of an action.

The mean expected value in the PEIRS model was also updated using Eq. (1). A measure of the spread in reward outcomes was learned in an analogous manner to Q-values using:

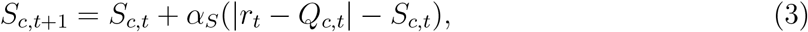

where is the learning rate for the spread, and *r_t_* − *Q_c,t_* is the reward prediction error that captures the deviation of the current outcome from the average outcome, which is compared with the current expected spread in reward outcomes *S_c,t_*.

The PEIRS model accounts for how participants differentiate matched mean stimuli based on the spread and captures individual risk propensities. For this model, the utility that was entered into the softmax function, Eq. (2), depends on the expected mean reward, the spread in reward outcomes and the sensitivity to the decision context (i.e. the context effect), in the following way:

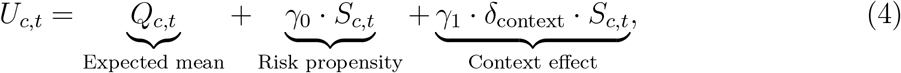

where the parameter *γ*_0_ modulates the risk propensity of an individual and reflects the tonic level of dopamine (Mikhael and Bogacz, 2016). A positive value of *γ*_0_ increases risk-seeking, because a high variance contributes positively to an option’s value, meaning that the high-spread option is preferred. This effect is reversed when *γ*_0_ < 0. Note that the first two terms in Eq. (4) are analogous to the mean-variance models developed for decisions from description (Boorman and Sallet, 2009; D’Acremont et al., 2009).

The third term captures the biasing effect of positive or negative decision contexts on choice behaviour. Context effects play an important modulatory role in risky decision-making (De Martino et al., 2006; Tversky and Kahneman, 1981) and were also observed in the current study. The context reflects how the expected value of the presented stimuli compares to the overall expected value of all stimuli in the task *δ*_context_ = *Q*_presented,t_ − *Q*_all,t_, where *Q*_presented,t_ is the average of the Q-values of the stimuli on the current trial and *Q*_all,t_ is the average of the Q-values of all four stimuli. The true average value of all stimuli is 50 points, but *Q*_all,t_ fluctuates around 50 as the Q-values of the stimuli change by trial-to-trial updates. Positive decision contexts have an objective value above the average 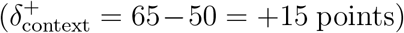, whereas negative decision contexts have a context value below the average 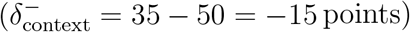. The parameter *γ*_1_ is a gain parameter that determines the extent to which the decision context and spread in reward outcomes contribute to choice behaviour. Positive values of *γ*_1_ increase risk-taking behaviour in positive contexts, and reduce risk-seeking in negative contexts. The opposite is true for negative values of *γ*_1_. In the PEIRS model, the effects of food deprivation can be attributed to how participants learn about the expected value (reflected by *α_Q_*), the spread of reward outcomes (reflected by *α_S_*), the individual risk propensity (reflected by *γ*_0_) and/or sensitivity to the context (reflected by *γ*_1_).

### Computational modelling captured risk preferences

A model comparison revealed that over 70% (23 out of 32 participants) were better described by the PEIRS model (BIC_RW_ = 16915 and BIC_PEIRS_ = 15996), confirming that the addition of extra parameters was justified. The quality of the fitting procedure was verified with a parameter recovery analysis. All parameters were well recovered (0.75 < *R* < 0.95) and the model fitting procedure did not introduce spurious correlations between the other parameters (|*R*| < 0.3; Fig. S4). Surrogate data generated with the best fitted parameters specifically confirmed that the model reproduces the key effect of food deprivation on choice preferences (Fig. 4A).

**Figure 4:**
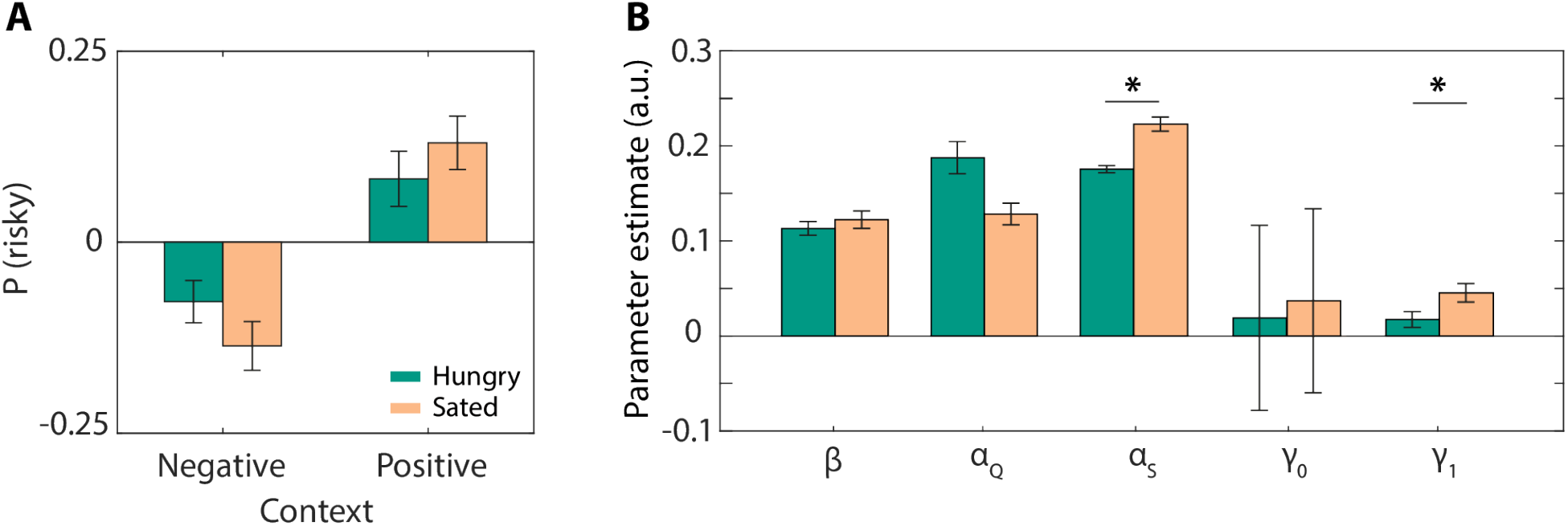
Model fitting results with the PEIRS model. **A)** Simulated choice behaviour using estimated parameters for the food deprived and sated condition. Simulated data showed a similar pattern to the behavioural data depicted in Fig. 3A. **B)** Food deprivation significantly decreased the learning rate for reward spread *α_s_*, and the sensitivity to contexts *γ*_1_. Food deprivation did not alter the softmax temperature *β*, the learning rate for mean *α_Q_* or individual risk preferences *γ*_0_. Error bars represent SEM. Statistical significance was tested with respect to the Gaussian scaled parameters. * *p* < 0.05.

In line with the behavioural analyses, we found an effect of food deprivation on parameter estimates obtained with the PEIRS model (Fig. 4B). On average, food deprived participants had lower learning rates for the spread [*α_S_*, *p* < 0.0001, *d* = 0.70] and a lower sensitivity to context effects [*γ*_1_, *p* = 0.02, *d* = 0.55], making them risk neutral across decision contexts. Food deprivation did not significantly alter learning rates of mean values [*α_Q_*, *p* = 0.165, *d* = 0.48] or choice stochasticity [*β*, *p* =1, *d* = 0.13]. Although risk propensities were differently affected by hunger among individuals, at the group level, individual risk propensities were not significantly altered by food deprivation [*γ*_0_, *p* = 1, *d* = 0.04].

### Subjective rating reflects learned utility

We also asked participants to indicate the reward distribution of each stimulus on a Visual Analogue Scale at the end of each block. We used these measures to examine whether people distinguished the stimuli based on the true mean and variance, or a scaled version of the objective values. We found that the subjectively rated mean and spread of each reward distribution (Fig. 5A–B) showed a similar pattern as the objective values (Fig. 1B). Participants were able to reliably distinguish stimuli based on their mean (main effect of mean [*F*_1,31_ = 831.91, *p* < 0.0001, 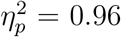]; Fig. 5A). The average outcome for the risky option was rated higher for high mean stimulus, but lower for the low mean stimulus (interaction effect of true mean and spread [*F*_1,31_ = 11.19, *p* < 0.002, 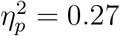]). To highlight this effect, Fig. 5A includes lines connecting the mean ratings of low and high variance stimuli, which have different slopes for high mean and low mean stimuli. These findings are consistent with the risk preferences observed in Fig. 3A, showing that participants valued their preferred stimulus more. This effect was less strong in hungry individuals, who rated the mean of stimuli in the positive and negative context more similarly, regardless of the level of risk (interaction effect of hunger and spread [*F*_1,31_ = 9.48, *p* < 0.004, 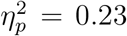]). The rated values are in line with the risk-neutral choice behaviour of hungry individuals (Fig. 3A).

**Figure 5:**
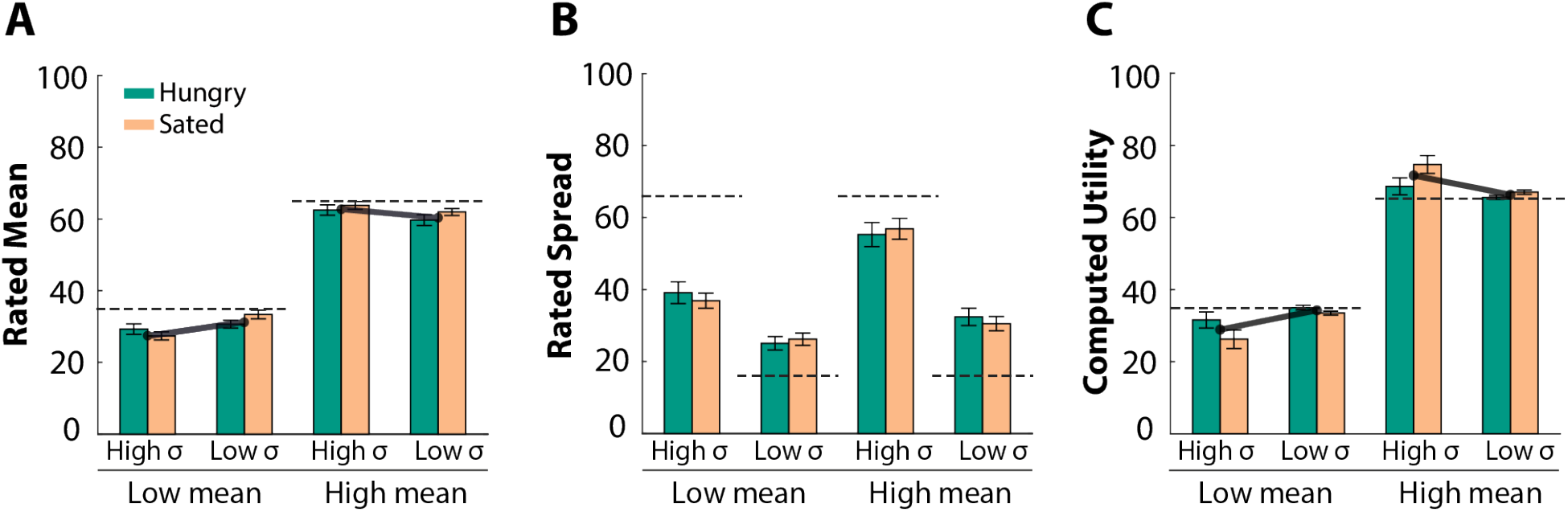
Subjective rating reflects learned utility. At the end of each block, participants indicated the mean **(A)** and spread **(B)** of the distribution associated with each stimulus. **C)** The computed utility for each of the stimuli, Eq. (4), reflects the same pattern as the subjectively rated mean values. Dashed lines indicate objective mean or spread in the reward points. Error bars represent SEM.

All participants understood that matched mean stimuli differed in the level of spread (main effect of spread [*F*_1,31_ = 61.93, *p* < 0.0001]; Fig. 5B). However, participants rated the spread for high mean options consistently higher than for low mean options (main effect of mean [*F*_1,31_= 86.56, *p* < 0.0001]). Furthermore, the perceived contrast in variance for high mean options was larger compared to the perceived contrast for low mean options (interaction effect mean and spread [*F*_1,31_ = 32.45, *p* < 0.0001, 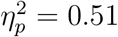]).

Given that we observed biases for the preferred (i.e. most chosen) stimulus in the subjective ratings, we examined whether this was reflected by the learned utility. The utility of each of the stimuli, Eq. (4), was computed using the *Q* and *S*-values obtained from the simulations in Fig. 4A and the best fitted parameters of each individual. During the ratings only one stimulus is presented at the time, thus *δ*_context_ = *Q*_rated stimulus_ − mean(*Q*_all stimuli_). The utilities were computed for each stimulus set separately and averaged across individuals (Fig. 5C). We observed three analogous effects in the learned utility as observed in the subjectively rated mean values (Fig. 5A vs Fig. 5C). First, the utility of high mean stimuli was significantly higher than the utility of low mean stimuli (main effect of mean [*F*_1,31_ = 319.85, *p* < 0.0001, 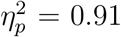]). Second, the learned utility for the risky option was higher for the high mean stimulus, but lower for the low mean stimulus (interaction effect of mean and variance [*F*_1,31_ = 19.32, *p* < 0.0001, 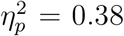]). Third, hunger altered the learned utility. Hunger increased the utility for low mean stimuli, but not for high mean stimuli (interaction effect of mean and hunger [*F*_1,31_ = 6.21, *p* = 0.018, 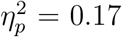]). This effect was specific for high variance options, but not low variance options (interaction effect of mean, variance and hunger [*F*_1,31_ = 5.86, *p* = 0.022, 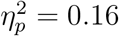]).

## Discussion

Using information about the current metabolic state to adapt to variable reward outcomes is critical for survival (Stephens, 1981). In this study, we used two complementary tasks to test whether food deprivation selectively affected risk-taking for learned or explicitly described options. We found that food deprivation modulated risk attitudes for decisions whose outcome statistics had to be learned, but not for decisions whose outcome statistics were explicitly described. Furthermore, hunger promoted risk aversion for positive contexts, but not for negative contexts. These results suggest that the current metabolic state drives adaptive behaviour for trial-and-error learning in a context-specific manner, but may not alter the integration of factual information.

As postulated by the risk-sensitive foraging theory (Stephens, 1981), individuals should make decisions that minimise the disparity between the goal and the current state to maximise the chance of survival. When forced to choose between two low reward options of similar expected value but different risk, the high variance option should be preferred when hungry, because this is the only option that offers a chance of fulfilling the current biological need (Fig. 6A). In contrast, when higher rewards are at stake, hungry individuals should now opt for the low-risk option, because this option allows them to fulfil their need, without incurring an unnecessary cost that may compromise survival (Fig. 6B). Although the participants in this study were not starving and the rewards in this task may only indirectly (via money) fulfil their biological needs, we found shifts in risk preferences (Fig. 3A) that follow the predictions by the risk foraging theory explained in Fig. 6. In particular, when participants became hungry, they decreased the tendency of selecting risky option in positive context (Fig. 3A). Our data illustrates that hunger has the tendency to alter risk-taking and provides evidence of how evolutionary pressure from the past is still influencing our behaviour.

**Figure 6:**
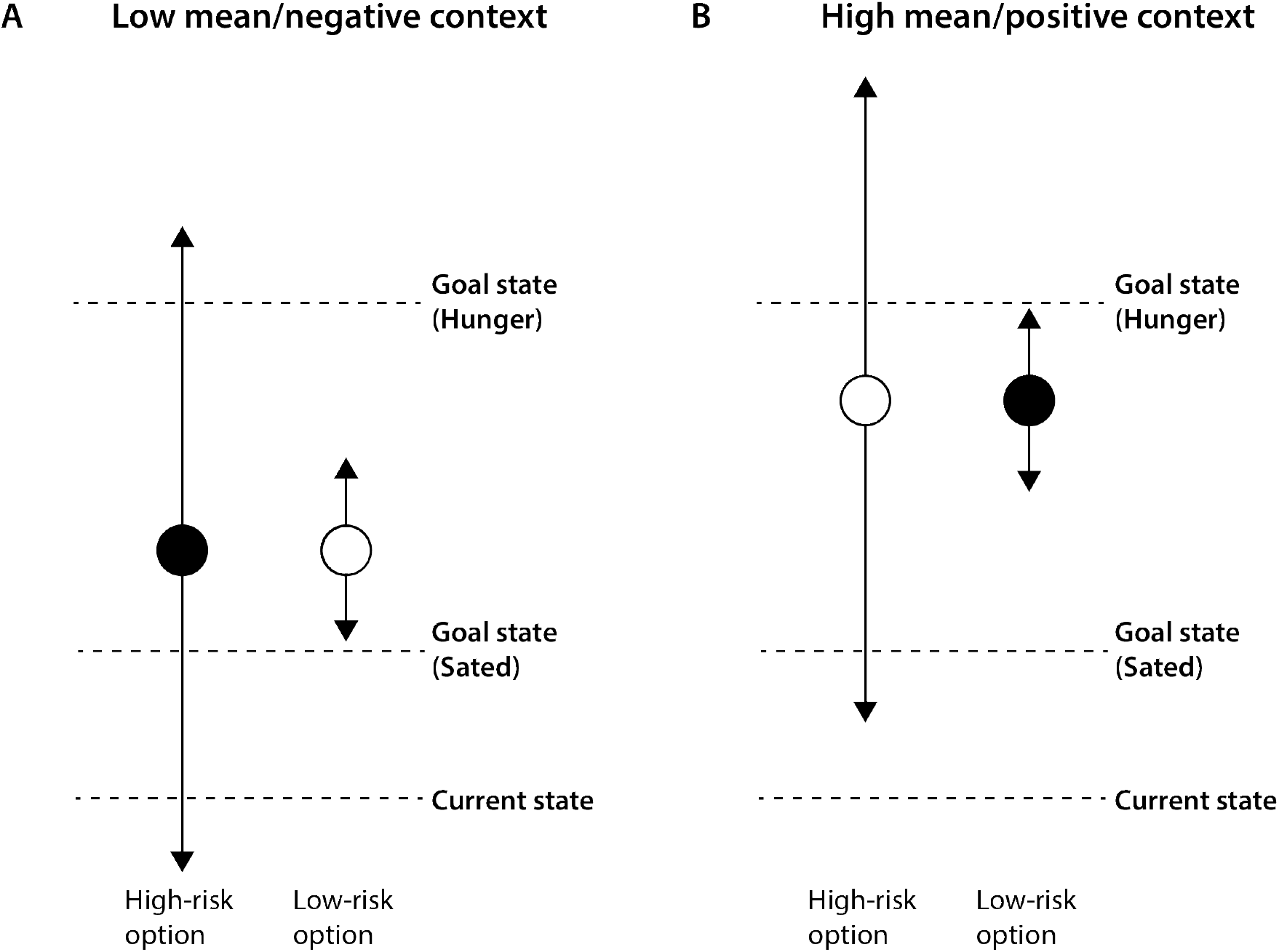
“Optimal” choice scenarios for positive and negative decision contexts. The circles denote the expected value of the high and low risk option and the arrows denote the spread of the reward. Filled circle indicates the preferred choice. **A)** Represents a scenario in a low mean/negative decision context. When forced to choose between options of similar expected value but different risk (i.e. outcome variance), decision-makers should prefer high-risk options (filled circle) when hungry (because it is the only option that offers a chance of fulfilling their need), and prefer low-risk options (open circle) when sated to ensure the goal state is achieved and avoid unnecessary downside costs that might be incurred if the high risk option is chosen. **B)** Represents a scenario in a high mean/positive decision context. The goal state can now be achieved with the low-risk option so this should be chosen in a positive context. The risky option should only be chosen if the needs cannot be met by choosing a safe option. Sated individuals can afford the costs (as this is still close to their goal state) and may therefore be more willing to gamble.

While both explicit and experiential risk-taking are modulated by the contextual value of the options presented, they are not equally susceptible to modulation by hunger. For experience-based decisions, information about the availability of reward and the metabolic need is integrated (Abizaid et al., 2006; Aitken et al., 2016; Cone et al., 2016; Hommel et al., 2006; Papageorgiou et al., 2016), whereas the evaluation of description-based decisions is susceptible to reward availability only.

Importantly, the behavioural data showed that the decision context was important for choice behaviour. For example, participants preferred the risky option in positive decision contexts, but preferred the safer option when it was presented with a low mean stimulus in a mixed context. This contextual adaptability is beneficial for survival and recent work has provided a mechanistic explanation for these contextual effects in experiential risk-taking (Möller et al., 2021). Pupil dilation at the time of decision context tracked how surprising the context was, corresponding to |*δ*_context_|. Furthermore, across individuals this dilation independently correlated with the size of *γ*_1_, which controls how strongly context biases choices. Crucially, in the present study, the effects of hunger were directly reflected by this parameter. Sated individuals showed a different choice bias in each decision context, while hungry participants were risk-neutral across both decision contexts. Hunger has been previously associated with maladaptive behaviour (Bartholdy et al., 2016; Kirk and Logue, 1997; Skrynka and Vincent, 2019), however, the results in this study show that hunger makes people more “rational” in their behaviour. These individuals rely more on the objective expected value of an option, rather than the subjective expected utility (von Neumann and Morgenstern, 1944).

Hunger did not affect description-based risk-taking. This may be surprising given that previous studies reported that hungry individuals were more risk-seeking for food, water and monetary rewards when gambles were explicitly described (Levy et al., 2013; Shabat-Simon et al., 2018; Symmonds et al., 2010). The absence of an effect may be due to differences in task design. First, previous studies mostly concerned a decision between a fixed certain amount and a risky alternative (Levy et al., 2013; Shabat-Simon et al., 2018), whereas the current study compared two risky options (as in Symmonds et al. (2010)), so one possibility is that hunger affects how risk is compared against certainty. Second, our task included 10 unique choice types that were played 8 times each, which might increase familiarity and promote explicit rational processing; in contrast previous studies used trial-unique gambles that were only played once. Third, studies involving described risks typically omit feedback. Although this approach is acceptable for a laboratory setting, real world choices usually lead to outcomes even if the outcome probabilities are known. Feedback about described risks could alter risk attitudes (Jessup and O’Doherty, 2010), we did not find evidence that learning occurred in this task as there was no change in risk preferences over the course of the task or across sessions.

We first opted for a design that was more similar in reward outcomes to the experienced-based task (similar to the design by Symmonds et al. (2010)), but a pilot study showed that using normally distributed reward complicated the task and failed to induce clear risk preferences. We therefore opted for a task that has been previously used to measure changes in risk preferences following the manipulation of the dopamine (motivational) system (Norbury et al., 2013).

Our data suggests that hunger does not impact risk-taking for description-based choices, at least when explicitly comparing two risky options with feedback provided, perhaps because the neural processes that drive explicit risk-taking are not under the direct control of hunger.

An important contribution of the current study is that we compared the effect of hunger on risk preferences for description and experience-based risks in the same individual following the same level of deprivation. Existing studies may be difficult to compare due to varying levels of deprivation; some studies report 4 hours deprivation, others 12 hours (Levy et al., 2013; Shabat-Simon et al., 2018; Symmonds et al., 2010). Furthermore, risk preferences vary greatly among individuals (Levy et al., 2013). Previous studies showed that hunger had a converging effect on a population – individuals who were highly risk-averse when satiated became less averse when hungry, while risk-seeking individuals became more risk-averse (Levy et al., 2013).

Our study demonstrates that the opposing risk patterns in the description-experience gap are driven by how risks are presented, rather than individual risk propensities. Previous studies suggested that these different risk patterns arise from memory biases (Madan et al., 2014) or under-and overweighting of rare events in description and experience-based choices, respectively (Hertwig, 2012; Hertwig et al., 2004; Kahneman and Tversky, 1979). The dissociable effect of hunger on experiential and explicit risk-taking in this study suggest that the neural processes driving these preferences are, at least partially, distinct (Fitzgerald et al., 2010; Jessup and O’Doherty, 2010).

In conclusion, we found that food deprivation decreased risk-taking for positive decision context in decisions where outcome statistics had to be learned. This observation matches optimal foraging theory, which predicts a survival advantage when individuals consider the variability of resources in the environment according to the current level of energy reserves. For learned risks, hungry individuals considered their metabolic need and the availability of rewards when making choices, whereas sated individuals only considered the availability for rewards. Hunger did not alter explicit risk-taking, suggesting that cognitive evaluation of risk may be unaffected. This is the first study that uses a within-subject design to test the effects of food deprivation on risk attitudes for decisions involving learned and described risks in positive and negative decision contexts. It provides new insights into the modulatory role of hunger in adaptive behaviour. Further studies will need to address the neural processing that are involved in the effects of hunger on decision-making under uncertainty.

## Acknowledgements

This work was supported by Medical Research Council grants MC_ST_U16043, MC_UU_12024/5, MC_UU_00003/1, MR/P00878/X, and BBSRC grant BB/S006338/1.

