## Supplemental material for "Gambling on an empty stomach: Hunger modulates preferences for learned but not described risks"

### Learning occurred for experienced, but not for described risks

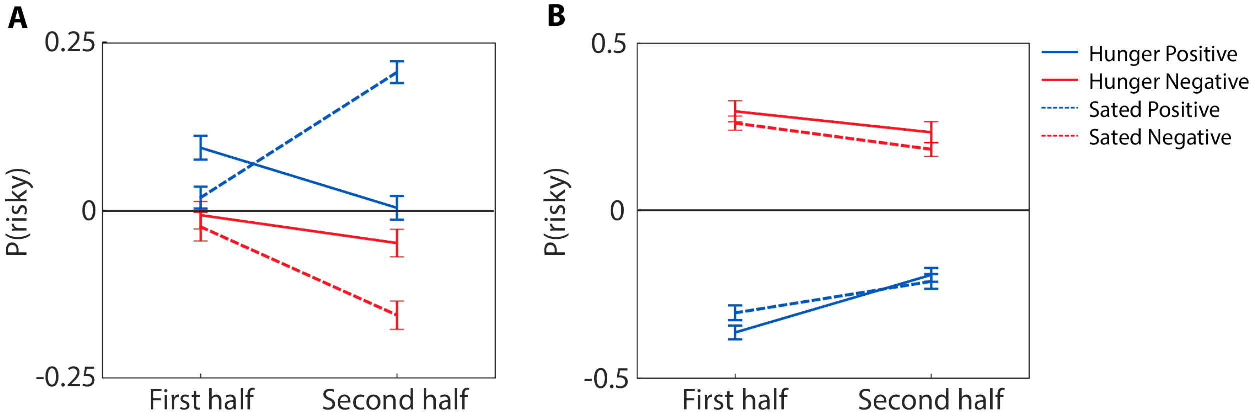


Figure S1: **Learning occurs in experienced-based risk-taking, not in description-based risk-taking.** Mean proportion of risky choices for the positive (blue) or negative (red) contexts as a function of blocks for the hungry (solid lines) and sated (dotted lines) condition. **A)** In the experience-based task, individuals are more risk-seeking for positive contexts than for negative contexts (main effect context [$F_{1,31} = 8.87$, $p < 0.006$, $\eta_p^2 = 0.22$]). Risk preferences change over time, but differently dependent on the decision context (interaction effect of time and context [$F_{1,31} = 6.69$, $p = 0.015$, $\eta_p^2 = 0.18$]), showing that learning occurs in this task. This effect is also modulated by the level of hunger (interaction effect of food deprivation, context and time [$F_{1,31} = 6.34$, $p = 0.017$, $\eta_p^2 = 0.17$]). **B)** In description-based risk taking, individuals are more risk-seeking for negative contexts compared to positive contexts (main effect context [$F_{1,31} = 55.01$, $p < 0.0001$, $\eta_p^2 = 0.64$]). Risk preferences do not change over time or as a result of food deprivation (all $p$-values $> 0.1$). Each bin contains equal number of positive and negative context trials. Error bars represent within-subject SEM.

### Pronounced context effects in a third of the participants

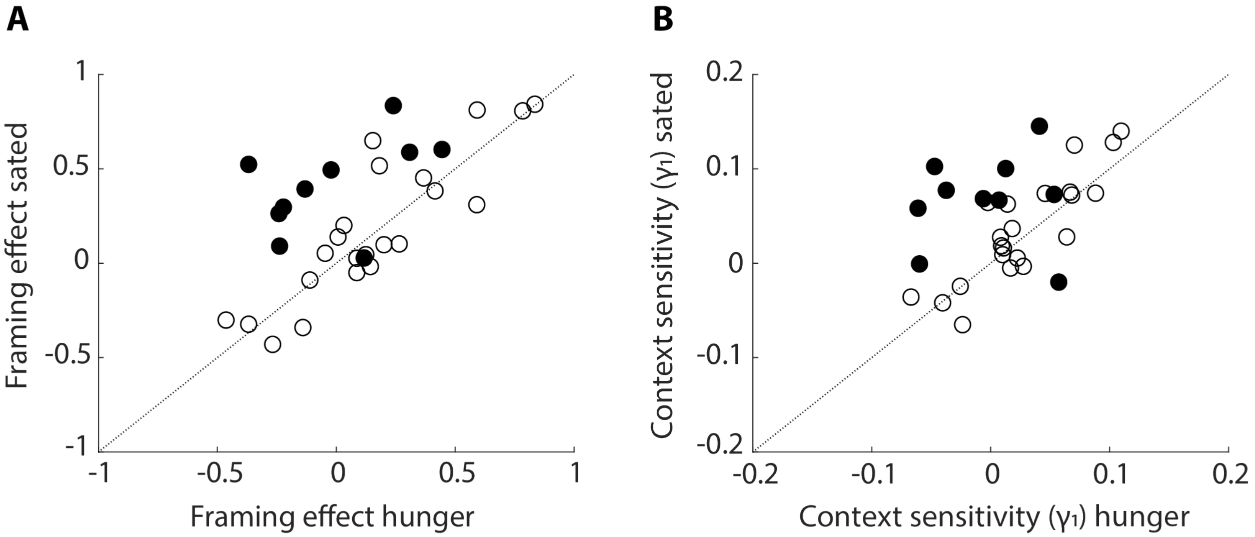


Figure S2: **Pronounced context effects in a third of the participants.** Participants with a significant interaction effect of context × hunger in the logistic regression are represented by filled circles. **A)** Individuals with a significant interaction effect show a greater framing effect when sated The framing effect is measured as P(risky | positive context) − P(risky | negative context). **B)** Individuals with a significant interaction effect showed greater context sensitivity ($\gamma_1$ parameter derived from computational modelling) when sated.

### Hunger increased reaction times for described, but not experienced risks

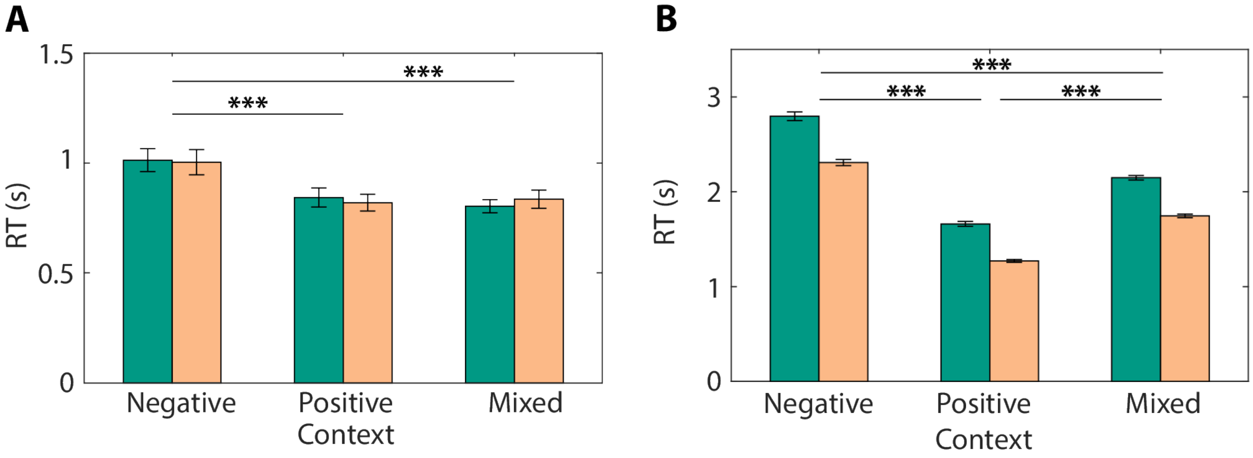


Figure S3: **Reaction times for for experienced and described risk.** **A)** Median reaction times for experience-based risk-taking, split by context type. Hunger did not affect overall reaction times for experienced risks (main effect of food deprivation [$F_{1,31} < 1$]; Suppl. Fig. S3A). Participants' reaction times did vary depending on the decision context (main effect of decision context [$F_{2,62} = 37.67$, $p < 0.0001$, $\eta_p^2 = 0.64$]). Further analyses using a two-way repeated measures ANOVA revealed that participants took significantly longer to respond for negative decision contexts compared to positive contexts [$F_{1,31} = 51.39$, $p < 0.0001$, $\eta_p^2 = 0.62$] and mixed contexts [$F_{1,31} = 39.19$, $p < 0.0001$, $\eta_p^2 = 0.56$], without showing a significant difference in reaction times between positive contexts and mixed contexts [$F_{1,31} = 0.55$, $p = 0.464$]. These data suggest that negative contexts were perceived as more difficult than positive or mixed contexts, a finding that has been reported before (Madan et al., 2015). Food deprivation did not affect context-specific reaction times (interaction effect of food deprivation and context [$F_{2,62} < 1$]). **B)** Median reaction times for description-based risk-taking, split by context type. Hunger increased reaction times for described risks (main effect of food deprivation [$F_{1,31} = 37.42$, $p < 0.0001$, $\eta_p^2 = 0.31$]; Suppl. Fig. S3B). Participants took significantly longer to respond for negative decision contexts compared to positive contexts [$F_{1,31} = 61.96$, $p < 0.0001$, $\eta_p^2 = 0.67$] and mixed contexts [$F_{1,31} = 18.79$, $p < 0.0001$, $\eta_p^2 = 0.38$]. They also deliberated longer for choices in mixed decision contexts compared to positive contexts [$F_{1,31} = 25.80$, $p < 0.0001$, $\eta_p^2 = 0.45$]. Error bars represent SEM. *** $p < 0.001$.

### Computational modelling

#### Parameter transformations

Before fitting the parameters, we applied logistic/hyperbolic/exponential transformations to transform bounded parameters into normal distributed parameter values $x_i \sim \mathcal{N}(\mu_x, \sigma_x)$, with a population mean of $\mu_x$ and a standard deviation of $\sigma_x$. We transformed [0,1]- bounded $\alpha_Q$, and $\alpha_S$ into a Gaussian scale using the logistic function:

$$\alpha = 1/(1 + \exp(-a)), \tag{1}$$

transformed [-1,1] - bounded $\gamma_0$, and $\gamma_1$ into a Gaussian scale using a hyperbolic tangent:

$$\gamma = \frac{\exp(g) - \exp(-g)}{\exp(g) + \exp(-g)}, \tag{2}$$

and the logarithmically scaled $\beta$ using the exponential function:

$$\beta = \exp(b) \tag{3}$$

We denote the model parameters by Greek letters and the Gaussian transformation by their respective latin letters. Normally distributed parameters allow for the use of parametric tests to identify differences between conditions. The statistical significance was tested using paired t-tests with respect to the Gaussian scaled model parameters. P-values were corrected for multiple comparisons using the Bonferonni method.

#### Model comparison

To identify the best fitting model, we compared the models using the Bayesian Information Criterion (BIC) value (Schwarz, 1978), which considers differences in model complexity. The BIC value was calculated as:

$$BIC = 2L + k \ln n, \tag{4}$$

where $L$ is the negative log likelihood, $n$ the number of choices and $k$ the number of free model parameters. The BIC value was computed using the maximum likelihood function of the population data across conditions. The model with the lowest BIC values was the best fitting model.

#### Parameter recovery

To validate the parameter estimates generated by the fitting procedure, we conducted a parameter recovery analysis. For each parameter, we generated samples from the marginalised posterior distribution of the fit to get realistic parameters that could describe choice behaviour. The generated parameters were uncorrelated ($|R| < 0.3$), allowing for testing whether the fitting procedure introduced any confounding factors. We used the generated sets of parameters to simulate choice behaviour and used a hierarchical model fitting procedure to estimate parameters for the simulated data ("Recovered parameters"). We then assessed the quality of the parameter recovery by comparing the true parameters used to simulate data with the recovered parameters. We calculated the Pearson correlation between all pairs of recovered parameters to test whether the fitting procedure introduced spurious correlations. A strong correlation between the true and recovered parameters indicates a good recovery of the parameters and reliable model fitting results. The quality of the fitting procedure was verified with a parameter recovery analysis. All parameters were well recovered

($0.75 < R < 0.95$) and the model fitting procedure did not introduce spurious correlations between the other parameters ($|R| < 0.3$; Fig. S4).

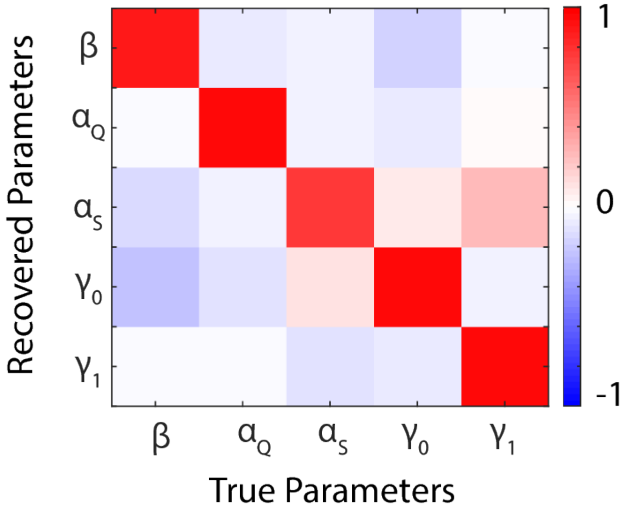


Figure S4: **Parameter recovery for the PEIRS model.** Correlation matrix of the free parameters used to generate the simulated data ('True parameters') and the obtained parameters by applying the parameter estimation procedure on the simulated data ('Recovered parameters'). Bright red values indicate a strong correlation between the true and recovered parameter value and therefore a good parameter recovery.
